# Development of a CMS Classifier for Clinical CRC Samples

**DOI:** 10.1101/2023.06.28.546869

**Authors:** Arezo Torang, Simone van de Weerd, Veerle van Lammers, Sander van Hooff, Jan Koster, Jan Paul Medema

## Abstract

Colorectal cancer (CRC) is a leading cause of cancer-related deaths worldwide, emphasizing the need for improved predictive biomarkers to guide treatment decisions. A classification system based on gene expression profiles, known as CMS, has shown promise in stratifying CRC into distinct subtypes with varying clinical outcomes. However, the lack of a reliable assay to classify formalin-fixed paraffin-embedded (FFPE) samples poses a challenge for translating this system into routine clinical practice. In this paper, we introduce the NanoClassifier, an NanoString-based CMS classifier to classify both FFPE and Fresh frozen (FF) tumors. We demonstrate the strong accuracy of the NanoClassifier in predicting CMS for CRC samples. By validating its performance on FF and FFPE samples, we highlight the prognostic significance of CMS in CRC.

## Introduction

Colorectal cancer (CRC) is a significant contributor to cancer-related mortality worldwide, ranking third in incidence^1–3^. While pathological staging is the current clinical practice for selecting patients for adjuvant chemotherapy^4^, it often fails to accurately predict recurrence in patients undergoing curative surgery for localized CRC, with around 25-35% of patients receiving adjuvant chemotherapy still experience the development of metastatic disease.

Furthermore, it is estimated that approximately 50-60% of these patients would have remained free of the disease even without undergoing adjuvant chemotherapy treatment^5,6^. Therefore, there is an urgent need for the development of additional prognostic and predictive biomarkers to improve patient selection for therapy with the aim to reduce unnecessary toxicities, and optimize treatment cost-effectiveness.

As a part of an international consortium, we previously proposed a classification scheme based on gene expression profiles from nearly 4000 high-quality RNA samples, dividing CRC into four biologically distinct subtypes; CMS1-4^7^. CMS1 mainly exhibits tumors with microsatellite instability (MSI), characterized by the presence of activated immune cells infiltrating the tumor. CMS2 and CMS3 both display epithelial features, with CMS2 primarily characterized by elevated WNT and MYC signaling, while CMS3 is marked by expression of genes regulating metabolic pathways. CMS4, on the other hand, comprises more mesenchymal-like cancers associated with significant stromal infiltration and poor patient outcomes^7^. While the classification system was primarily built using whole transcriptomic data from fresh-frozen samples, the lack of a suitable assay to classify patient samples into subtypes using formalin-fixed paraffin-embedded (FFPE) samples with clinically relevant turn-around times and costs poses a challenge in translating these findings into routine clinical practice.

In order to fully realize the potential of the CMS system, there is a need for a robust, reliable single-sample classifier that can efficiently process RNA extracted from FFPE samples. This paper introduces the NanoClassifier, an FFPE-based CMS classifier utilizing the NanoString platform, which demonstrates strong accuracy in predicting CMS for colorectal cancer samples. We validated the performance of this FFPE tissue-based gene classifier and demonstrated the prognostic significance of CMS in colorectal cancer, paving the way for improved patient outcomes and more targeted treatment strategies.

## Materials and Methods

### RNA Extraction

For RNA extraction from FF tissue, we followed the manufacturer’s protocol using RNA-B and Trizol. Exclusion criteria for samples were set as a tumor percentage of less than 30%. When working with FFPE tissue, we prepared three 10 μm slides that were subjected to H&E staining. De-paraffinisation of tissue was carried out using Xylene, followed by macro-dissection of the specimen to enrich for tumor tissue. RNA extraction was performed using the RNeasy FFPE kit (Qiagen, location) as per the manufacturer’s instructions. To assess the quality and quantity of RNA, we used the NanoDrop2000 and Tapestation (Agilent).

### RNAseq Profiling

Prior to read trimming and alignment, the quality of raw data was assessed using FastQC (v.0.11.9). Trimming was performed on 12 and 11 bases for single-end and pair-end RNAseq data, respectively. To further enhance data quality, bases with a Phred score less than 20 were filtered using Cutadapt (v1.18). The reads were aligned to the reference genome (GRCh38) and gene counts were generated using the STAR (v.2.7.4.a) algorithm. To enable accurate downstream analyses, the count data was log2-transformed and quantile normalization was applied.

### NanoString Profiling

To quantify the expression levels of genes of interest in both FF and FFPE samples, we utilized 50 - 150 ng of total RNA and two custom-designed CodeSets. Additionally, we included both positive and negative probes to monitor the quality of runs. To ensure reliable data analysis, we performed log2-transformation and quantile normalization as our standard method of normalization in this study.

### CRC Gene Expression Datasets

In this study, we utilized 12 publicly available datasets (GSE13067, GSE13294, GSE14333, GSE17536, GSE20916, GSE2109, GSE23878, GSE33113, GSE35896, GSE37892, GSE39582, TCGA^8^) and two proprietary datasets (PETACC3, KFSYSCC) which were made available through Synapse (www.synapse.org) (ID: syn2623706). PETACC3 was derived from FFPE (n = 688) and we obtained normalized data from each study.

### CodeSet1

For the custom nCounter assay dedicated to CMS subtypes, an initial selection of 134 genes was made based on differential expression analysis on the TCGA dataset.

This initial custom CodeSet1 was employed to profile training samples obtained from FF materials (AMC90-FF, n = 41; Biobank-FF, n = 21), and when feasible, from their corresponding FFPE tissues (AMC90-FFPE, n = 33; Biobank-FFPE, n = 16), utilizing the NanoString platform, which were named FF-NanoString dataset (n = 62) and FFPE-NanoString dataset (n = 49), respectively. For all these samples whole transcriptomic data of FF tissues was available (GSE103340, GSE33113), and the CMS labels were determined as the gold standard by employing the SSP function in the CMSclassifier package^7^.

### CodeSet2

In an effort to minimize the number of genes in the CodeSet1 as much as possible without compromising accuracy, we utilized the feature selection capability of the glmnet package^9^. For FF materials, we first merged the FF-NanoString dataset with TCGA (n = 570) and GSE39582 (n = 585), ensuring each sample from TCGA and GSE39582 was first quantile normalized by NanoString samples only using the genes in CodeSet1. Subsequently, 80% of FF-NanoString dataset were randomly selected and, in conjunction with TCGA and GSE39582, were used to train a model and test the remaining 20% to check the accuracy. This process was repeated 1000 times, each time accumulating the selected genes. Following this, the same procedure was implemented, but with the updated list of genes, and it was continued until there was no drop in accuracy. As a result, a total of 51 genes were identified to define the CodeSet2.

The CodeSet2 was then employed to profile a set of test samples (MATCH-FF, n = 165; DELFT-FFPE, n = 33) in order to validate the classifiers. In parallel, RNAseq data of FF tissues was generated for all DELFT samples (n = 33) and MATCH samples with sufficient RNA quality (n = 113) to determine the gold standard CMS labels by utilizing the SSP function in the CMSclassifier package^7^.

### Construction of a FF-NanoClassifier for CMS

We employed a weighted domain adaptation approach to leverage the FF-NanoString dataset and also microarray and RNAseq datasets as training sets. To train a FF-NanoClassifier which faithfully classifies fresh frozen materials, we utilized a combined set of CMS-stratified TCGA and GSE39582 samples (n = 923) in addition to FF-NanoString dataset (n = 62), using the genes in CodeSet2. Each sample of TCGA and GSE39582 was first individually quantile normalized with FF-NanoString dataset to align with the same distribution (Figure 1). After normalization of the training set, we employed Elastic-net, a logistic regression model, using cv.glmnet function implemented in the glmnet R package^9^. The selected Lasso coefficient was 0.7, and weights for NanoString samples were optimized to 1.5 while this number for RNAseq and microarray samples was set to 1.

**Figure 1.**
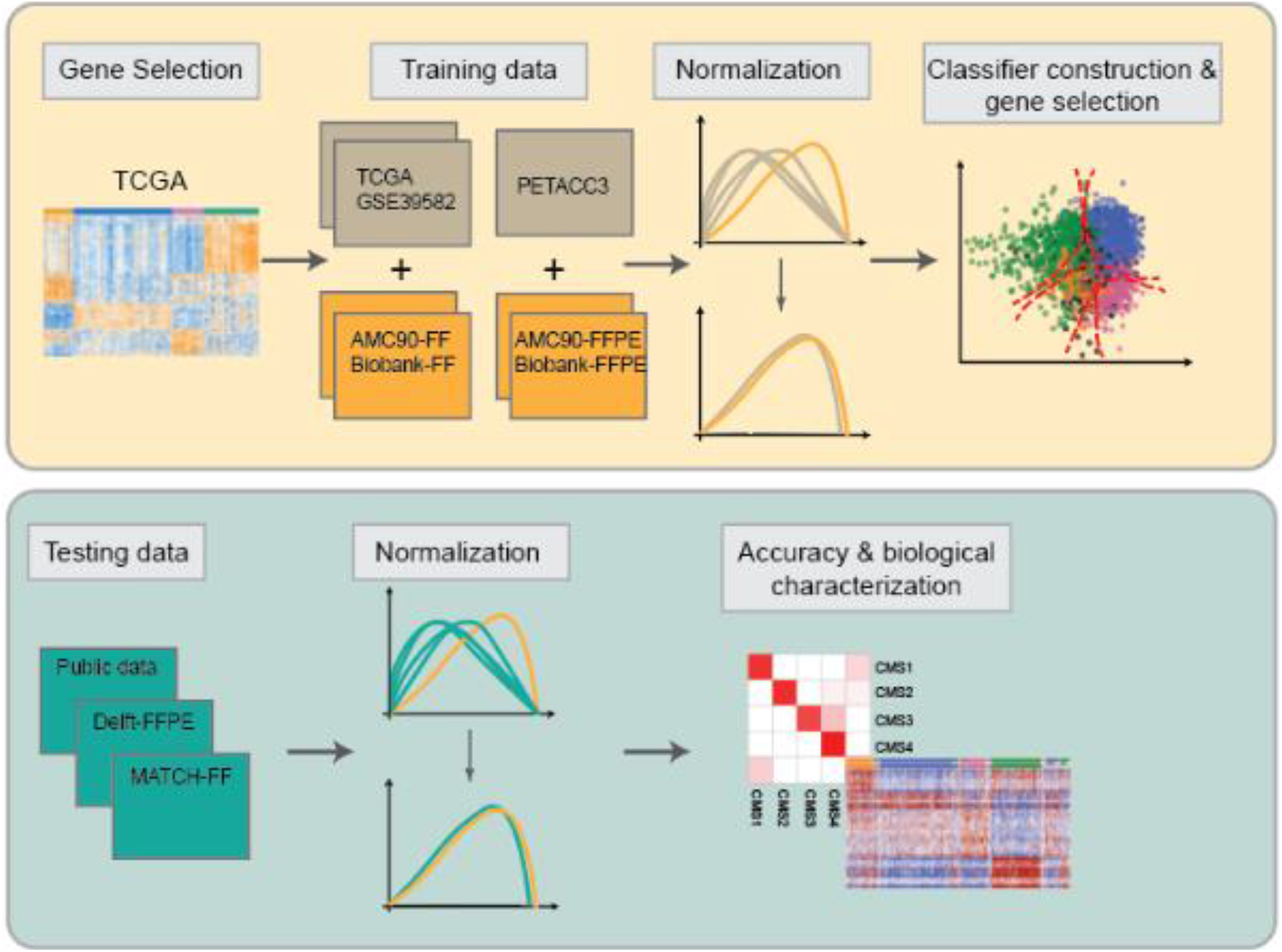
Workflow of training and testing the NanoClassifier.

### Construction of a FFPE-NanoClassifier for CMS

For the FFPE-NanoClassifier, the CMS-stratified PETACC3 samples (n = 537) were used in conjunction with FFPE-NanoString dataset (n = 49) to normalize the PETACC3 samples, using the genes in CodeSet2. Each PETACC3 sample was first quantile normalized with FFPE-NanoString dataset to align with the same distribution. Subsequently, 80% of FFPE-NanoString dataset were randomly selected and, in conjunction with PETACC3, were used to train a model and test the remaining 20% of FFPE-NanoString data to obtain the accuracy. This process was repeated 1000 times, each time accumulating the selected genes. Following this, the same procedure was implemented, but with the updated list of genes, and it was continued until there was no drop in accuracy. This procedure was done to avoid overfitting and as a result 41 genes were obtained to develop the FFPE-NanoClassifier. Using this 41 genes, the same normalization method was repeated on PETACC3 samples. Next an Elastic-net model was used to train the classifier. The selected Lasso coefficient for this model was 0.8, and weights for NanoString samples were optimized to 1.5 while this number for RNAseq and microarray samples was set to 1. The workflow is shown in Figure 1.

### Single Sample Gene Set Enrichment Analysis

To obtain pathways and signatures that are specific to each CMS subtype, we extracted data from MSigDB (v7.4) or Synapse (www.synapse.org) (ID: syn2623706). To calculate the scores of these signatures for each sample, we implemented the Z-score method of the gsva function (GSVA package^10^).

### Principal Component Analysis

In order to perform Principal Component Analysis (PCA) on the public CRC datasets, we initially employed the normalize.quantiles function of the preprocessCore package to normalize the collected samples. Subsequently, we utilized the prcomp function from the stats package^11^ to conduct PCA.

## Results

### The Selection of Genes for NanoClassifier Development

To facilitate the subtyping of colorectal cancer samples obtained from archival clinical material, we developed a customized gene-expression panel using NanoString technology. The NanoString platform is a well-established method for profiling RNA derived from formalin-fixed paraffin-embedded (FFPE) samples^12,13^. Leveraging the TCGA data, we identified genes with discriminatory power in delineating CMS subtypes through differential expression analysis. The CMS labels of TCGA samples were determined using the SSP classifier. Figure 2A illustrates the differential expression of these selected genes across various CMS subtypes in datasets profiled using different platforms, demonstrating that this expression pattern remains consistent regardless of the transcriptomic profiling platform employed. Despite the notable decline in gene expression quality observed in FFPE materials, the distinguishing pattern remained discernible.

**Figure 2.**
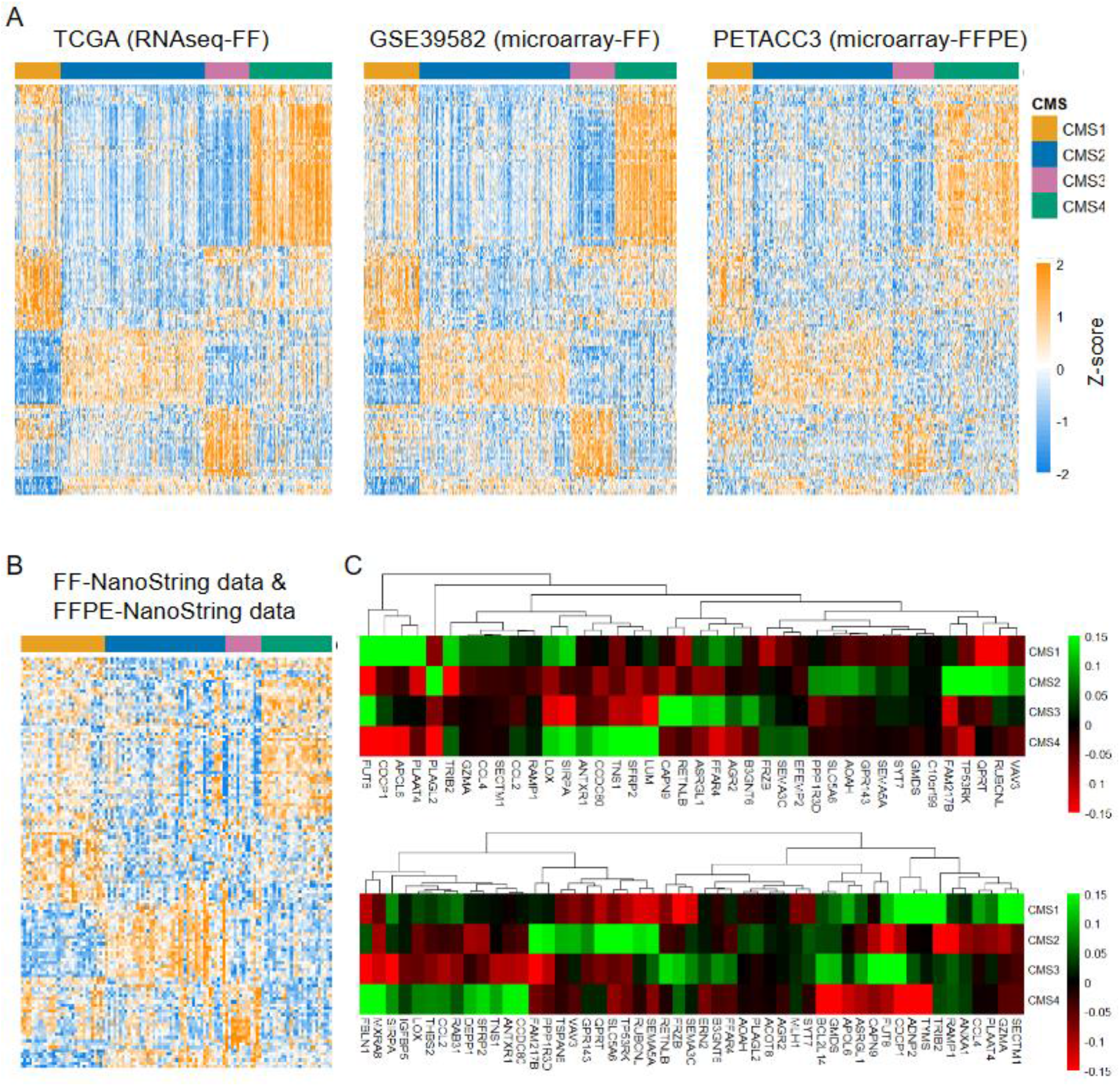
Visualization of CodeSet1 and NanoClassifier Coefficients. A. Heatmap of genes in CodeSet1 for whole transcriptomic datasets with defined CMS, TCGA (n = 457), GSE39582 (n = 466), PETACC3 (n = 537). B. Heatmap of genes in CodeSet1 for combined FF-NanoString and FFPE-NanoString datasets C. Coefficients of final selected genes in FFPE-NanoClassifier (top) and FF-NanoClassifier (bottom)

To further evaluate the suitability of this gene list for the NanoString platform, we visualized the combined FF-NanoString data and FFPE-NanoString data in Figure 2B. Once again, the pattern effectively captured the CMS subtypes in these samples. It is important to note that all CMS labels in this figure were obtained using the SSP classifier. In order to enhance cost-efficiency, this gene set was further condensed to a minimal number of 51 (CodeSet2), as explained in the materials and methods section.

### Development of FF and FFPE-NanoClassifier

To create tissue-specific classifiers capable of accurately classifying FF or FFPE materials, we developed a NanoString-based classifier using CodeSet2. For training the FF-NanoClassifier, we utilized a dataset comprising CMS-stratified TCGA and GSE39582 samples (n = 923), along with the FF-NanoString dataset (n = 62), and employed the Elastic-Net model. A similar approach was employed for the FFPE-classifier, using CMS-stratified PETACC3 samples (n = 537) alongside the FFPE-NanoString dataset (n = 49). However, the number of genes in the FFPE-classifier model was reduced to 41 genes. This reduction was implemented to prevent overfitting, which can compromise the generalization of the model^14^ (See the materials and methods section for more details). Figure 2C displays the coefficients of the genes in both models.

### NanoClassifier for RNAseq and Microarray Data

To assess the performance of our classifier, we tested it on a set of 2738 samples from publicly available datasets with CMS labels obtained from the Synapse^7^ as the “gold-standard”. As shown in Figure 3A a high concordance of 95% between the “gold-standard” labels and our NanoClassifier predictions was observed, with over 86% of samples being confidently classified (probability score > 0.6). Moreover, we conducted PCA on the samples and the first two principal components were visualized in Figure 3B. It was observed that the unclassified samples in both SSP and NanoClassifier tended to cluster at the boundaries between CMS groups, where CMSs intersect, which indicated that these samples were difficult to classify.

**Figure 3.**
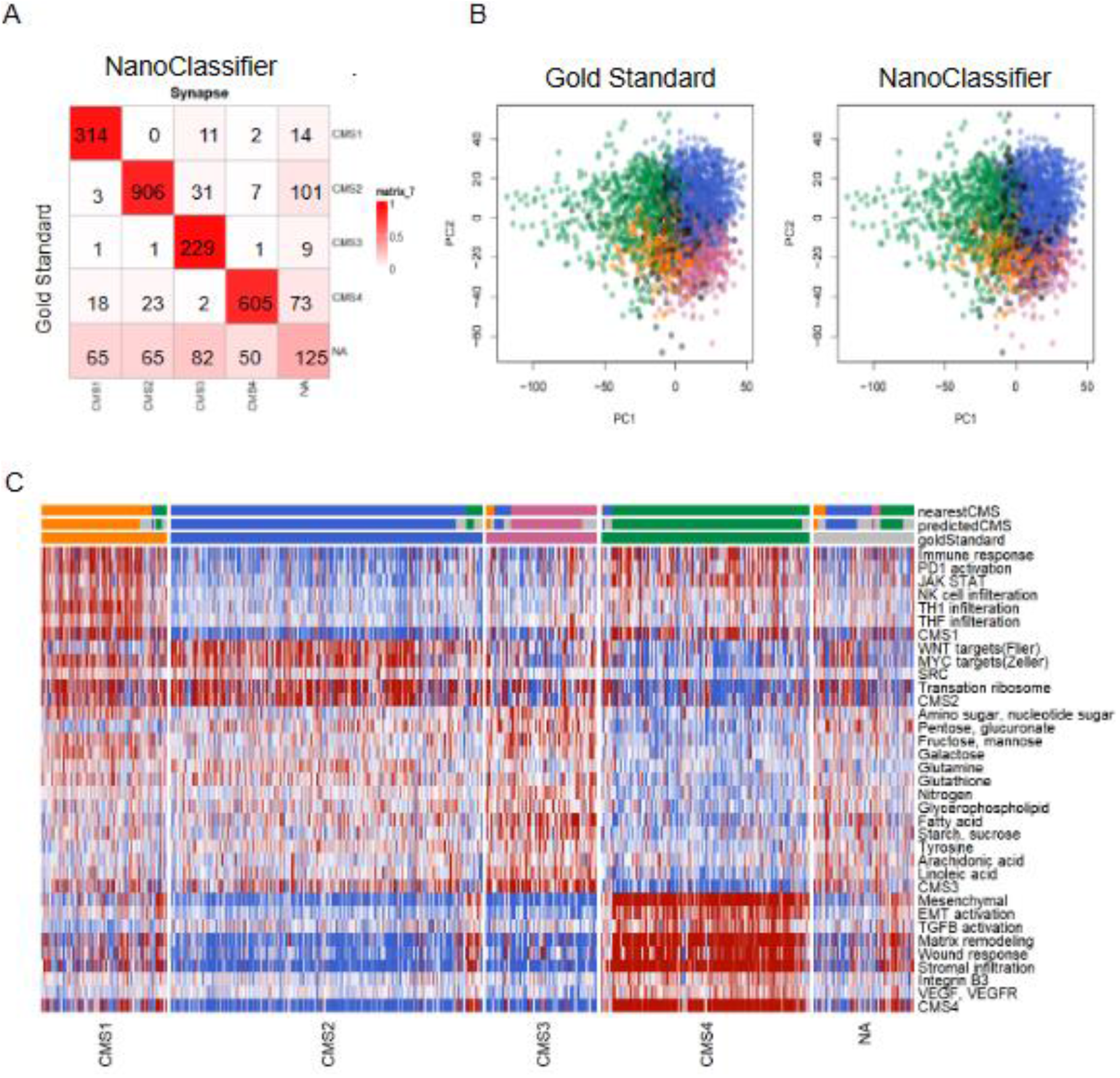
Classification of Whole transcriptomic Data. D. Concordance matrix of the results of SSP and NanoClassifier stratification of combined public data (n = 2738). E. PCA plot of combined public data colored by CMS labels of SSP classifier (left) and NanoClassifier (right) F. ssGSEA heatmap of combined public data, samples are clustered by the SSP stratification.

### Biological Characteristics of CMSs Were Retained by NanoClassifier

To further evaluate the ability of our NanoClassifier to faithfully stratify tumour samples with the proper biological characteristics^7^, we scored for the presence of CMS-specific pathways and signatures for each sample. As depicted in Figure 3C, confidently classified samples exhibited a clear enrichment of the expected CMS-specific signatures. This confirmed that even a small set of genes present in our NanoClassifier could accurately stratify samples based on the CMS system. Notably, samples that did not meet the defined probability threshold had weaker signals for almost all signatures, indicating a lack of enrichment for any of the pathways of interest.

Collectively, our findings highlight the ability of the NanoClassifier to accurately predict CMS using RNAseq and microarray data. Moreover, our results suggested that samples with low probability scores are less likely to exhibit the biological characteristics defining CMSs.

### NanoClassifier for FF and FFPE - NanoString Datasets

To further evaluate the performance of the NanoClassifier, we applied it to two new sets comprising 113 FF (MATCH-FF) and 33 FFPE (DELFT-FFPE) samples, which were profiled using the newly developed CodeSet2 on the NanoString platform. As RNAseq was available for these samples as well the “gold-standard” CMS labels were obtained using the SSP classifier from the CMSclassifier package^7^. Comparing the NanoString-based analysis combined with the NanoClassifier labels with the gold-standard labels revealed an accuracy of 95% using FF-NanoClassifier for MATCH-FF and 92% implementing FFPE-NanoClassifier for DELFT-FFPE (Figure 4). To our knowledge, this represents the highest accuracy reported thus far for CMS stratification on FFPE material^15–18^. These results demonstrated the superior performance of our NanoClassifier and its ability to accurately stratify FF and FFPE samples. Especially the latter is a crucial step towards clinical implementation as FFPE material is standardly acquired in clinical routine.

**Figure 4.**
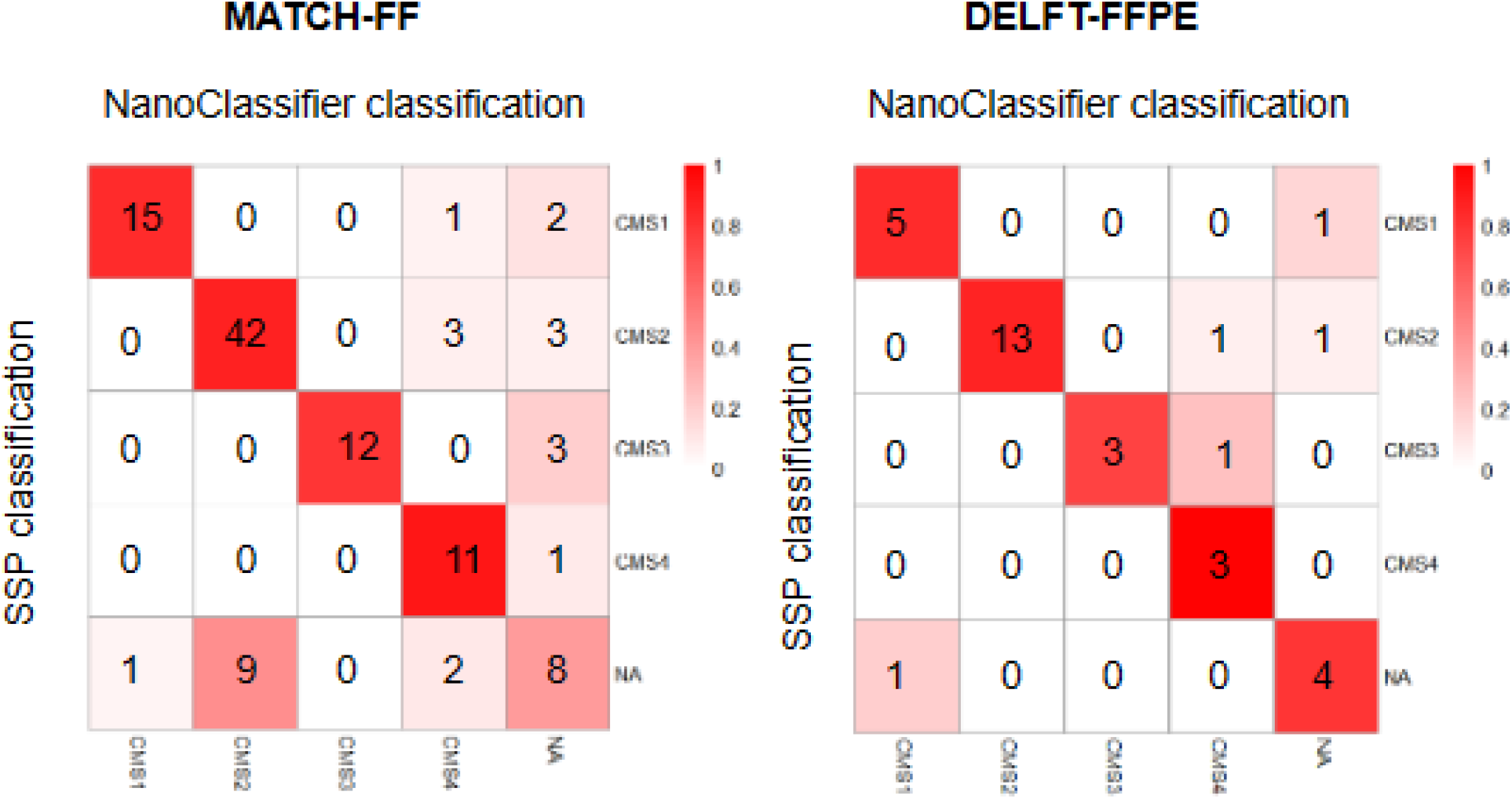
Classification of Whole transcriptomic Data. Concordance matrix of the results of SSP and NanoClassifier stratification of MATCH-FF (left; n = 113) and DELFT-FFPE (right; n = 33)

## Discussion

The development of accurate and reliable prognostic and predictive biomarkers is crucial for improving patient outcome and optimizing treatment strategies in colorectal cancer (CRC). In this study, we introduced the NanoClassifier, a NanoString-based CMS classifier for FF/FFPE tissues, and demonstrated its strong accuracy in predicting CMS for CRC samples.

The CMS classification system has emerged as a valuable tool for stratifying CRC into biologically distinct subtypes with varying clinical characteristics and outcomes^19^. However, its widespread adoption in clinical settings has been hindered by the lack of a suitable assay for classifying FFPE samples, which are commonly used in routine pathological practice. Our study addressed this challenge by developing the NanoClassifier, which overcomes the limitations of existing methods and provides a robust single-sample classifier for FFPE, as well as, FF-based CMS classification.

Our findings demonstrated that the NanoClassifier confidently classified a significant proportion of samples, surpassing the performance of the SSP classifier. This outcome underscores the potential of the NanoClassifier to offer valuable information for making personalized treatment decisions in the context of CRC. Moreover, samples classified using the NanoClassifier showed a pronounced presence of CMS-specific pathways and signatures, highlighting the biological significance of the CMS classification system. This evidence bolsters the idea that CMS subtypes possess unique molecular features, and a precise categorization of CRC samples can yield important prognostic details. Earlier, it was proposed that understanding CMS could predict both the therapeutic response and the likelihood of disease recurrence, specifically peritoneal metastasis^20,21^. Moreover, the CMS4 subtype, known as the mesenchymal subtype, is characterized by significant stromal invasion with the worst survival^22^. Therefore, determining the CMS subtype would enable a more personalized therapeutic approach.

Previously, Morris et al. developed a NanoString-based classifier for archival material, achieving an 80% accuracy rate^16^. Although there are notable differences in accuracy rate and validation methods between their study and ours, both studies indicate the reliability of the NanoString platform in developing and validating gene expression-based signatures using FFPE samples. Furthermore, Ragulan et al. successfully utilized the NanoString platform to develop a classifier consisting of 38 genes for the CRCA system, a subtyping model they developed for colorectal cancer. Their classifier achieved high accuracy in CRCA classification, however, it yielded a lower accuracy rate of 75% when applied to the CMS subtyping system^18^. Similarly, Marisa et al. developed an RF classifier using the NanoString platform, but only 69% of the samples were assigned to a CMS subtype with a probability score of 0.5 or higher^17^.

While other studies have also utilized the NanoString platform, demonstrating its potential in developing gene expression-based classifiers, variations in accuracy rates and subtype assignment highlight the complexities involved in achieving accurate CMS classification for archival material. Nonetheless, these collective findings emphasize the value of the NanoString platform and its potential to improve our understanding of CRC and guide personalized treatment strategies.

In conclusion, our study presents the NanoClassifier as a robust and accurate FF and FFPE-based CMS classifier for CRC samples. By overcoming the limitations of existing methods, the NanoClassifier holds great potential for translating the CMS classification system into routine clinical practice, enabling more precise and targeted treatment strategies for CRC patients. The prognostic significance of CMS subtypes and the practical advantages of the NanoClassifier pave the way for improved patient outcomes.

